# Lipids with negative spontaneous curvature decrease the solubility of the cancer drug paclitaxel in liposomes

**DOI:** 10.1101/2023.10.18.563006

**Authors:** Victoria Steffes, Scott MacDonald, John Crowe, Meena Murali, Kai K. Ewert, Youli Li, Cyrus R. Safinya

## Abstract

Paclitaxel (PTX) is a hydrophobic small-molecule cancer drug that loads into the membrane (tail) region of lipid carriers such as liposomes and micelles. The development of improved lipid-based carriers of PTX is an important objective to generate chemotherapeutics with fewer side effects. The lipids 1,2-dioleoyl-*sn*-glycero-3-phosphoethanolamine (DOPE) and glyceryl monooleate (GMO) show propensity for fusion with other lipid membranes, which has led to their use in lipid vectors of nucleic acids. We hypothesized that DOPE and GMO could enhance PTX delivery to cells through a similar membrane fusion mechanism. As an important measure of drug carrier performance, we evaluated PTX solubility in cationic liposomes containing GMO or DOPE. Solubility was determined by time-dependent kinetic phase diagrams generated from direct observations of PTX crystal formation using differential-interference-contrast optical microscopy. Remarkably, PTX was much less soluble in these liposomes than in control cationic liposomes containing univalent cationic lipid 1,2-dioleoyl-3-trimethylammonium-propane (DOTAP) and 1,2-dioleoyl-*sn*-glycero-3-phosphatidylcholine (DOPC), which are not fusogenic. In particular, PTX was not substantially soluble in GMO-based cationic liposomes. The fusogenicity of DOPE and GMO is related to the negative spontaneous curvature of membranes containing these lipids, which drives formation of nonlamellar self-assembled phases (inverted hexagonal or gyroid cubic). We used synchrotron small-angle x-ray scattering to determine whether PTX solubility is governed by lipid membrane structure (condensed with DNA in pellet form) or by local intermolecular interactions. The results suggest that local intermolecular interactions are of greater importance and that the negative spontaneous curvature-inducing lipids DOPE and GMO are not suitable components of lipid carriers for PTX delivery regardless of carrier structure.

## Introduction

Paclitaxel (PTX) [1] is one of the most commonly used cancer chemotherapy drugs, with applications in the treatment of ovarian, breast, lung, and other cancers [2-7]. Because PTX is a prototypical hydrophobic drug with poor water solubility, it has to be delivered using a carrier. In most clinical applications, patients receive PTX intravenously in the form of Taxol^®^ [8], a formulation of PTX dissolved in a solvent mixture of polyethoxylated castor oil and ethanol which has undesired clinical side effects [9-12]. More recently, the PTX formulation Abraxane^®^ was approved, which solubilizes PTX in the nontoxic and biocompatible albumin protein in nanoparticle form [13-17]. However, this second generation treatment is still dose-limited by tolerability of the PTX side effects and has not definitively improved patient outcomes compared to Taxol. To overcome the serious side effects of PTX-containing therapeutics stemming from the PTX carriers as well as from off-target effects of PTX against healthy cells, new delivery systems for PTX and other cytotoxic drugs are being developed [12,13,18-24].

Liposomes are versatile drug carriers that are used in approved therapeutics and in clinical trials worldwide [19,25-34]. They can be loaded with diverse cargo: hydrophilic small molecule drugs can be encapsulated in their interior, or liposomes can be complexed with nucleic acids (DNA or RNA) to form lipid–nucleic acid complexes (lipoplexes) or lipid nanoparticles (LNPs) [35-38]. Of particular relevance to the present study, hydrophobic drugs such as PTX can also be loaded into liposomes. These compounds intercalate into the membranes of liposomes to be solubilized by the fatty acid tails [19,39-41] (with the exception of few specific compounds that allow loading of crystalline material into in the aqueous core by unique mechanisms [42-44]). PTX delivery efficacy (i.e., cytotoxicity against cancer cells) correlates with the extent of time that PTX remains soluble in the lipid membranes after hydration [39]. The prolonged drug solubility likely keeps PTX bioavailable during the period between carrier administration and the time PTX reaches its biomolecular target, tubulin, in the cytoplasm. The mechanism of action of PTX involves binding to the β-tubulin subunit of microtubules, which results in mitotic block and apoptosis [3,45-50]. Therefore, we have endeavored to develop lipid-based carriers of PTX with long-term membrane solubility of PTX at high membrane concentrations. This will minimize side effects by requiring less carrier and because the high efficacy of the carrier will allow administration of lower total doses of PTX, reducing off-target effects [35,39,51-54].

Cationic lipids are an essential component of the lipid nanoparticles (LNPs) recently developed for siRNA and mRNA delivery in therapeutics and vaccines [36-38,55-59] because of their ability to complex with anionic nucleic acids and achieve loading. When used in the delivery of small-molecule drugs, cationic liposomes are able to passively target solid tumors through electrostatic interactions [32,52,60-66]. Newly forming vasculature associated with growing tumors is more negatively charged than other tissue in the body, thereby attracting positively charged particles at higher concentrations. For this reason, some liposomal formulations for drug delivery also include cationic lipids. One example is EndoTAG^®^-1, a PTX formulation currently in clinical trials [12,62]. EndoTAG-1 consists of cationic 1,2-dioleoyl-3-trimethylammonium-propane (DOTAP), neutral 1,2-dioleoyl-*sn*-glycero-3-phosphocholine (DOPC), and PTX in a 50:47:3 molar ratio [19,27,29,30,67,68].

As part of the ongoing effort to develop improved lipid-based carriers of PTX, we sought to investigate whether LNP design principles would carry over to PTX delivery. For example, cholesterol is a major component (≈40 mol% of the lipid mixture) in the LNP formulations of the approved siRNA therapeutic OnPattro™ (patisiran) [55] and the mRNA vaccines against SARS-CoV-2 [56-59] and is frequently used as a major component in liposomal formulations for the delivery of *hydrophilic* drugs. Cholesterol increases liposome bending rigidity in membranes in the lipid chain melted state and also increases the permeability barrier, thus reducing loss of the drug from diffusion across the membrane [42,69,70]. However, cholesterol has been shown to compete with PTX for space in the membrane, ultimately decreasing PTX solubility and loading [19,40,41,71-73]. Therefore, cholesterol should not make up a large fraction of lipid-based formulations for PTX delivery.

Another common component of LNPs and lipople**x**es are lipids with a propensity to form membra es with negative spontaneous curvature (*C*_0_ < 0). These lipids are sometimes labeled “fusogenic” because the elastic properties of their membranes facilitate fusion with other lipid membra es. This has been postulated to help the membranes of lipoplexes and LNPs merge with endosomal membranes, facilitating delivery of their cargo into the cytoplasm [74-84]. Lipids promoting *C*_0_ < 0 have an inverted-cone molecular shape, with a lipid head group smaller than the tail area (Fig. 1). The two examples of such lipids which we used in this work are glyceryl monooleate (GMO [85-87] and 1,2-dioleoyl-*sn*-glycero-3-phosphoethanolamine (DOPE). DOPE is chemically similar to DOPC, but DOPC’s bulkier choline headgroup leads to a cylindrical molecular shape that promotes *C*_0_ = 0 [83]. Fig. 1 shows the molecular shapes, chemical structures, and the self-assembled morphologies of their lipoplexes with DNA for the lipids mentioned above.

**Fig. 1.**
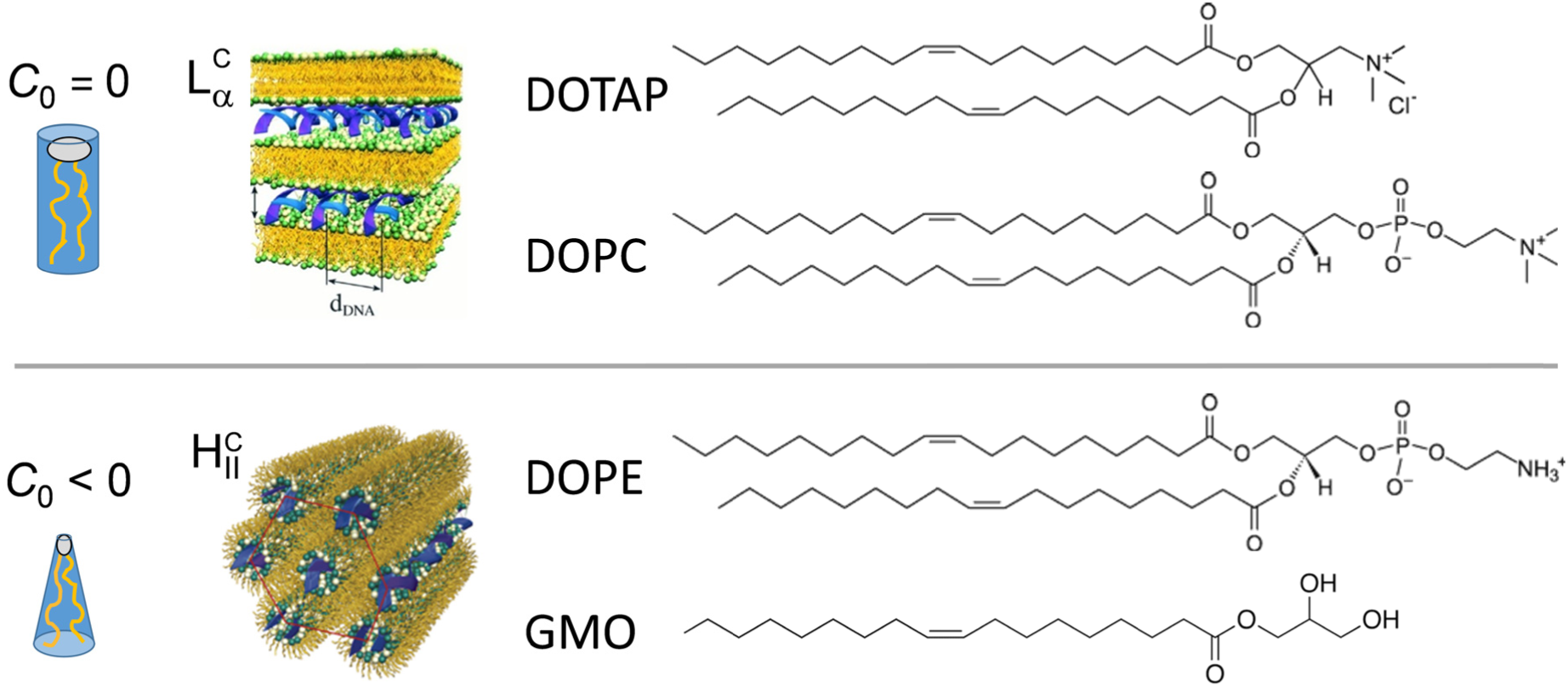
Molecular shape, self-assembled structures of lipoplexes, and chemical structures of the lipids used in this work. Lipids with cylindrical shape such as DOTAP and DOPC favor self-assembly into lamellar bilayer struct res and form lipoplexes in the L_α_^C^ phase upon condensation with DNA. Lipids with an inverted cone shape, such as DOPE and GMO, self-assemble into nonbilayer structures and yield related lipoplex structures such as the inverted hexagonal H_II_ phase. Schematics of lipoplex structures lipids (with lipid tails in yellow, headgroups in green and white, DNA as blue helices) adapted from [74]. Reprinted with permission from AAAS.

To investigate whether DOPE and GMO are suitable lipids for PTX delivery when compared to DOPC, we used differential-interference-contrast (DIC) microscopy to assess PTX solubility over time, generating kinetic phase diagrams for several lipid formulations. These formulations included binary lipid mixtures of cationic DOTAP with a neutral lipid (DOPC, DOPE, or GMO) as well as ternary lipid formulations of DOTAP with a mixture of DOPC and DOPE at various ratios. The PTX loading of the formulations varied between 0.5 and 5 mol%. Consistent with previous reports, we found that PTX remains solubilized for 24 hours or longer up to a content of about 3 mol% in formulations largely composed of DOPC [19,39-41]. Remarkably, PTX was significantly less soluble in formulations containing DOPE or GMO than in those containing DOPC, with solubility decreasing as the content of lipid favoring negative spontaneous curvature increased. This finding indicates that lipids favoring negative spontaneous curvature are ill-suited for PTX delivery.

To determine whether the decrease of PTX solubility is directly related to the liposome morphology, we performed small-angle x-ray scattering (SAXS) experiments on lipoplexes of the formulations where we complexed cationic liposomes with DNA. This allowed us to condense the liposomes into pellets suitable for quantitative structure determination. While we observed the expected inverse hexagonal (H_II_^C^) phase at high content of DOPE or GMO, the lamellar (L ^C^) phase and mixtures of the two phases were also observed at lower content of these lipids. Considering that PTX solubility was poor in the formulations independent of the lipoplex phase they formed, we conclude that lipid–PTX intermolecular interactions for DOPE and GMO have a stronger effect on PTX solubility than the liposome morphology.

## Materials and Methods

### Materials

Lipid stock solutions of DOTAP and DOPC in chloroform were purchased from Avanti Polar Lipids at 25 mg/mL (35.8 and 31.8 mM, respectively). DOPE and GMO were purchased as powders and dissolved in chloroform to a concentration of 50 mM. PTX was purchased from Acros Organics and dissolved in chloroform at a concentration of 10.0 mM. Calf thymus DNA was purchased from Thermo Scientific and dissolved in high-resistivity water (18.2 MΩcm) to a concentration of 3.5 mg/mL.

### Liposome Preparation

Solutions of lipid and PTX were prepared in chloroform/methanol (3:1, v/v) in small glass vials at a total molar concentration (lipid+PTX) of 5 mM for DIC microscopy and 30 mM for x-ray scattering experiments. Individual stock solutions of the components were combined according to the final desired molar composition and at the desired final concentration. The organic solvent was evaporated by a stream of nitrogen for 10 min and dried further in a vacuum (rotary vane pump) for 16 h. The resulting lipid/PTX films were hydrated with high-resistivity water (18.2 MΩcm) to the concentrations noted above.

### Kinetic phase diagrams

Following hydration, samples prepared at 5 mM concentration were mixed either by manual agitation or using a vortex mixer. The sample solutions were stored at room temperature for the duration of the experiment. At predetermined times, aliquots of 2 μL were withdrawn, placed on microscope slides, covered by a coverslip kept in place by vacuum grease, and imaged at 10× or 20× magnification on an inverted Diaphot 300 (Nikon) microscope. The samples were imaged every 2 h up to 12 h, then daily up to 10 days, and periodically thereafter until PTX crystals were observed or the sample was used up. The kinetic phase diagrams report the median time to observation of PTX crystals after hydration for 3–6 independently hydrated samples at each PTX content.

### Small angle x-ray scattering

To make x-ray samples, a total of 50 μL of a 30 mM liposome solution in water were mixed by vortexing in a 500 μL centrifuge tube with the volume of a solution of calf thymus DNA (3.5 mg/mL) required to achieve a cationic lipid to anionic DNA charge ratio of 1:1. The samples were subsequently centrifuged in a table-top centrifuge at 5,000 rpm, and the resulting pellets were transferred into 1.5 mm quartz capillaries (Hilgenberg) with excess supernatant. The capillaries were centrifuged in a capillary rotor in a Universal 320R centrifuge (Hettich) at 10,000 *g* and 25 °C for 30 min. After centrifugation, capillaries were sealed with two-component epoxy resin.

SAXS measurements were carried out at the Stanford Synchrotron Radiation Lightsource (Palo Alto, CA), beamline 4-2 at 9 keV (*λ*=1.3776 Å) with a Si(111) monochromator. Scattering data was measured by a 2D area detector (MarUSA) with a sample-to-detector distance of ≈3.5 m (calibrated with a control sample of silver behenate). The x-ray beam size at the sample was 150 μm in the vertical and 200 μm in the horizontal directions. Scattering data is reported as the radial average of scattering intensity against the scattering vector *q*.

## Results

### Optical microscopy to assess PTX solubility and phase separation

We used differential-interference-contrast (DIC) microscopy to monitor PTX-loaded lipid samples starting 2 hours (h) after hydration. Liposomes formed spontaneously upon hydration and were not sonicated. Figure 2 shows representative images of liposomes loaded with varying amounts of PTX (1.5, 2, 3, and 5 mol% across the horizontal axis). In addition to PTX, the lipid mixtures contained 70 mol% neutral lipid (mixtures of DOPC and DOPE, with the ratio of these lipids shown along the vertical axis) and the remainder cationic lipid (DOTAP). The formulation at 70 mol% DOPC shows PTX solubility and efficacy similar to the formulation of EndoTAG-1 with 50 mol% DOPC [73] and serves as our reference formulation.

**Fig. 2.**
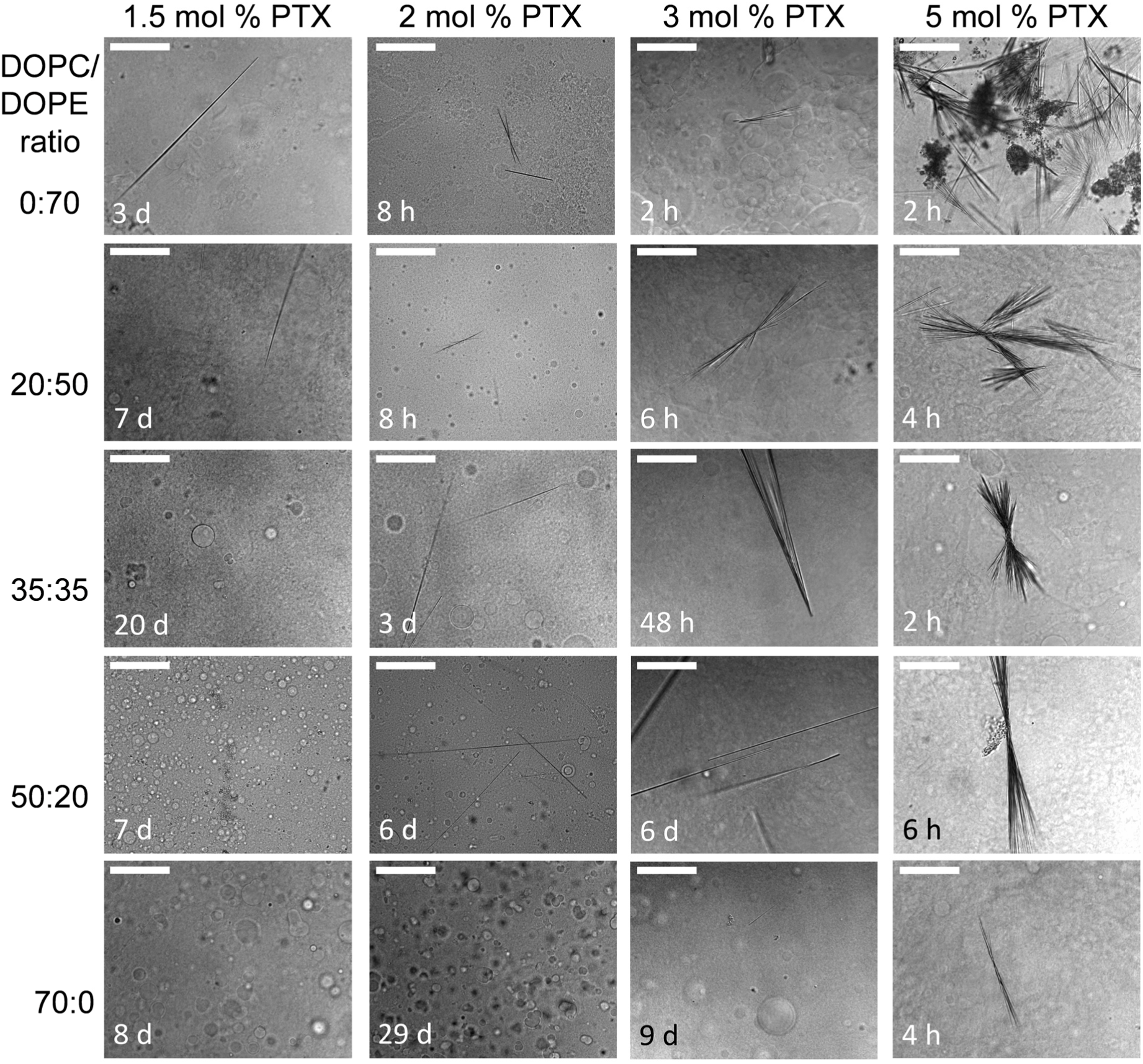
Representative DIC micrographs of cationic liposomes and phase-separated PTX. Lipid samples composed of DOPC:DOPE (total 70 mol%) at varying molar ratio (as indicated along the vertical axis), PTX (1.5–5 mol%; specified along the horizontal axis), and DOTAP (remainder) were monitored by DIC microscopy over time until PTX crystals were observed or until 20 days (d) after liposome hydration if no crystallization occurred. The time after hydrati**o**n at which the micrograph was taken is shown on each image. The appearance of the PTX crystals varied by composition. The time to PTX crystallization was extended at low PTX loading and at higher DOPC content; in other words, PTX crystallizaed more quickly from liposomes with high PTX loading and high DOPE content, often forming bundles of crystals. Scale bars: 100 μm.

The micrographs in Fig. 2 were selected to show PTX crystals unless the sample did not show any PTX crystallization within the window of observation of 20 days (d). Two main trends are evident: (i) the extent of crystallization was greatest and occurred most quickly for samples with the highest PTX loading, as expected; (ii) similarly, increasing DOPE content appeared to promote PTX crystallization. We often observed the formation of bundles of needle-shaped PTX crystals at high PTX loading and high DOPE content, whereas at lower PTX loading or higher DOPC content samples, individual PTX needles were more common.

### Kinetic phase diagrams: PTX solubility in binary lipid formulations

Plotting the data obtained by DIC microscopy as a function of PTX loading and time after to PTX crystallization hydration generates kinetic phase diagrams for the studied lipid formulation. Figure 3 displays these diagrams for lipid formulations containing PTX and binary lipid mixtures of cationic DOTAP mixed with a neutral lipid (DOPC, DOPE or GMO). The blue color indicates time points where PTX remained soluble, while the red color indicates the presence of phase separated PTX crystals. (The plotted data reflect the median time to PTX crystallization from three or more trials.)

**Fig. 3.**
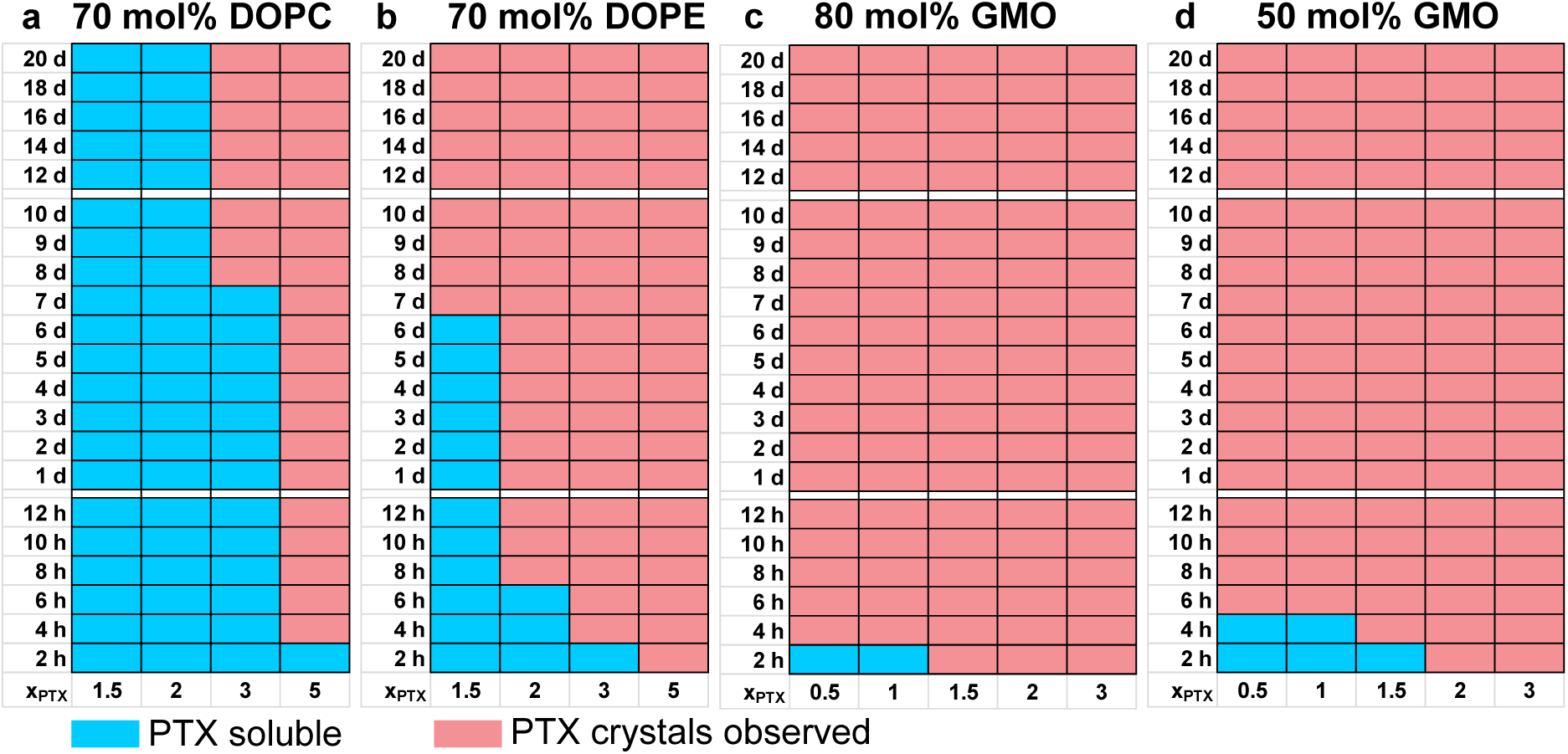
Kinetic phase diagrams of PTX solubility in binary lipid formulations of cationic DOTAP with neutral DOPC, DOPE, or GMO lipid. The blue color indicates that PTX remained completely solubilized by the lipid, while the red color indicates presence of PTX crystals. The border between the red and blue regions is the PTX membrane solubility boundary. The diagrams indicate the duration of PTX solubility in unsonicated liposomes containing a) 70 mol% DOPC, b) 70 mol% DOPE, c) 80 mol% GMO, and d) 50 mol% GMO. PTX was incorporated at 0.5–5 mol% of the formulations, as indicated along the x-axis, while the remaining lipid was cationic DOTAP. The solubility boundary was determined from the median of 3–5 separate trials at each PTX content.

The DOPC-based formulation, with a spontaneous curvature near zero (*C*_0_ ≈ 0), solubilizes PTX best, with no PTX crystals observed at 3 mol% PTX until 8 days after hydration (Fig. 3a). PTX solubility significantly decreases when DOPC is replaced with DOPE (with *C*_0_ < 0 because of the smaller size of the phosphoethanolamine headgroup compared to phosphocholine). Even at a content as low as 1.5 mol%, PTX is only soluble up to 6 days, while at 3 mol% PTX, crystals were observed 4 h after hydration (Fig. 3b).

To further explore the role of *C*_0_ and membrane morphology on PTX solubility, we also studied formulations with GMO as the neutral lipid. GMO has only a single lipid tail, but also a much smaller (glycerol) headgroup than DOPC or DOPE, resulting in a negative *C*_0_. GMO has shown the ability to form both inverted hexagonal and cubic self-assemblies [75,77,86-88]. Here we investigated the solubility of PTX in formulations with 50 and 80 mol% GMO. Initial experiments immediately indicated, both by microscopy and visible solution cloudiness, that PTX phase-separates rapidly from GMO formulations. Accordingly, we assessed PTX solubility at lower drug loading than for the formulations containing DOPC or DOPE, down to 0.5 and 1.0 mol% (as indicated along the horizontal axis in Fig.s 3c,d). Even at this low loading, PTX is poorly solubilized by membranes containing GMO as a major component. Decreasing the GMO content from 80 to 50 mol%, thereby increasing the amount of DOTAP (with *C*_0_ ≈ 0), only elicited a minor improvement of PTX solubility.

### Kinetic phase diagrams describing PTX solubility in ternary lipid formulations

Expanding on the data for the binary lipid compositions, we assessed PTX solubility in a series of ternary lipid formulations containing a mixture of neutral DOPC and DOPE (at molar ratios of 20:50, 35:35, and 50:20) with cationic DOTAP. The PTX membrane solubility boundaries for these formulations are superimposed in Fig. 4 (including the data for 70 mol% DOPC and 70 mol% DOPE from Fig. 3a,b).

**Fig. 4.**
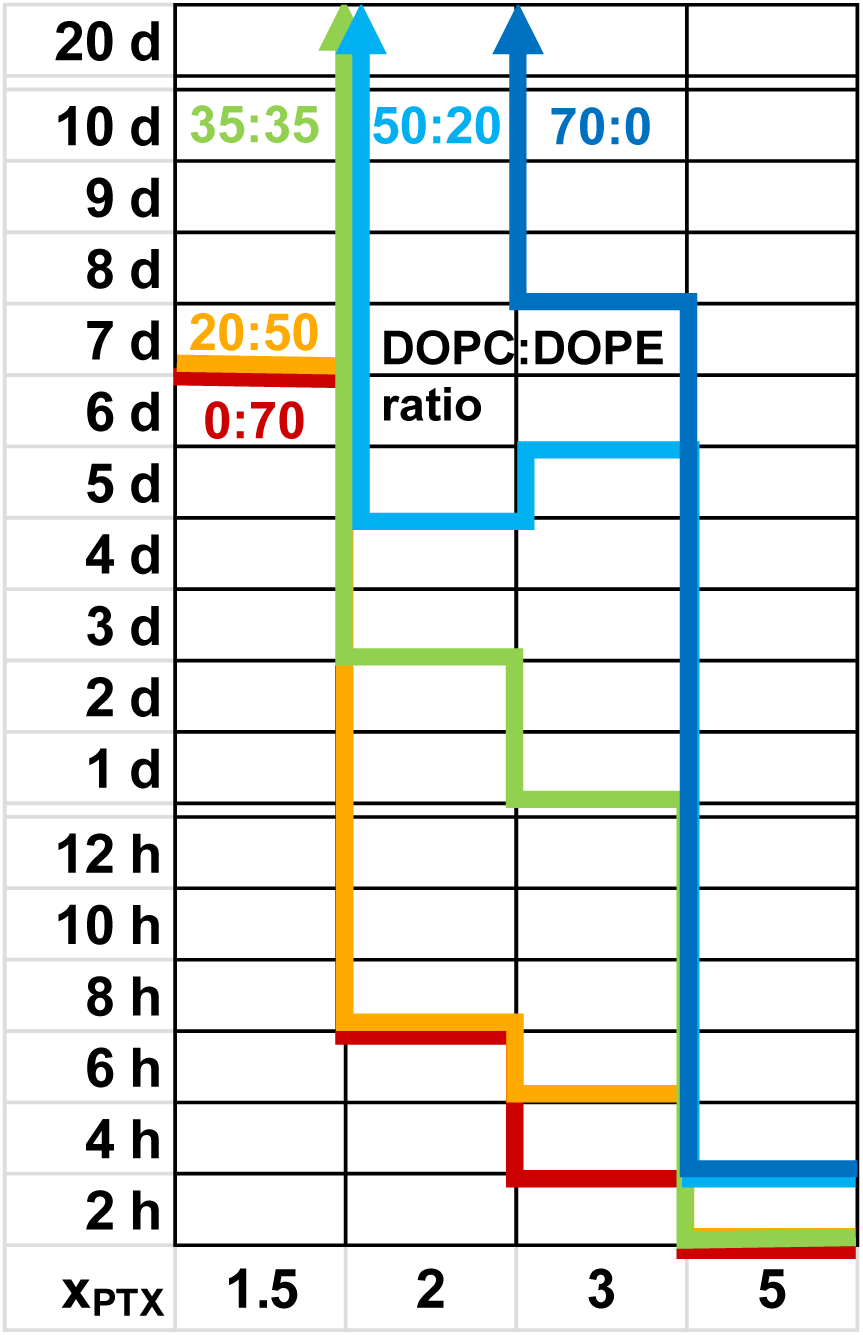
Solubility of PTX in ternary lipid formulations of cationic DOTAP with neutral DOPC and/or DOPE lipid. The figure shows an overlay of the PTX membrane solubility boundaries of kinetic phase diagrams obtained for five lipid formulations with 70 mol% neutral lipid at varied DOPC:DOPE ratio (0:70 in red; 20:50 in orange; 35:35 in green; 50:20 in light blue; 70:0 in dark blue). PTX content was 1.5–5 mol%, as indicated along the x-axis, and the remainder (25–28.5 mol%) was DOTAP. The PTX membrane solubility boundaries were determined from the median of 3–5 separate samples at each PTX content.

The formulations at DOPC:DOPE ratios of 0:70 and 20:50 (red, orange) show poor PTX solubility with nearly identical PTX membrane solubility boundaries. As the relative amount of DOPC increases, PTX solubility also increases. Even 20 d after hydration, no PTX crystals were observed at 1.5 mol% PTX loading for the formulations at of 35:35, 50:20 and 70:0 DOPC:DOPE (green, light blue, dark blue). At contents of 2 and 3 mol%, PTX was soluble in these formulations for at least a day. The formulation at 70 mol% DOPC was the only formulation where PTX was soluble for > 20 d at 2 mol% drug loading.

### Small angle x-ray scattering (SAXS) of condensed lipid-DNA samples

Because the shape results in a negative spontaneous curvature, both DOPE and GMO have a propensity for forming nonlamellar lyotropic phases and the inverted hexagonal phase in particular. The solubility experiments therefore raise the question whether membrane morphology is a factor affecting PTX membrane solubility. To elucidate this question, we used small-angle x-ray scattering (SAXS) to determine the self-assembly structures of cationic liposome-DNA complexes (i.e. lipoplexes) formed from a series of lipid formulations. These formulations contained 2 mol% PTX, neutral lipid as specified and the remainder DOTAP. we have previously performed SAXS experiments of lipoplexes prepared from PTX-loaded liposomes [39]. Electrostatically condensing the liposomes with DNA to form a pellet increases the coherent domain size and local concentration of material, thus increasing scattering and providing additional physical information that dilute solution scattering alone would not yield.

Fig.s 5 and 6 show the radially averaged SAXS intensity profiles for the lipoplexes. The SAXS intensity profiles in Fig. 5 are annotated with peak indexing for the lamellar (L_α_^C^) (00L) or inverse hexagonal (H_II_^C^) (HK) phases, as well as the visible DNA–DNA correlation peak of the L_α_^C^ phase [74,89,90]. The nanostructures of the L_α_^C^ and H_II_^C^ lipoplex phases are schematically shown in Fig. 1b. Of the four binary lipid formulations, those with 70 mol% DOPC and 50 mol% GMO yield lipoplexes with L_α_^C^ morphology, while the ones with 70 mol% DOPE and 80 mol% GMO yield lipoplexes in the H_II_^C^ phase. PTX crystallization results in characteristic peaks at *q*_P1_ = 0.291 Å^-1^, *q*_P2_ = 0.373 Å^-1^, and *q*_P3_ = 0.436 Å^-1^, consistent with our previous report [39]. These peaks are visible in all of the samples except for the one prepared at 70 mol% DOPC, consistent with the solubility studies outlined in the previous section. The data show that PTX crystallizes faster from liposomes and lipoplexes containing DOPE and GMO than from those containing DOPC.

**Fig. 5.**
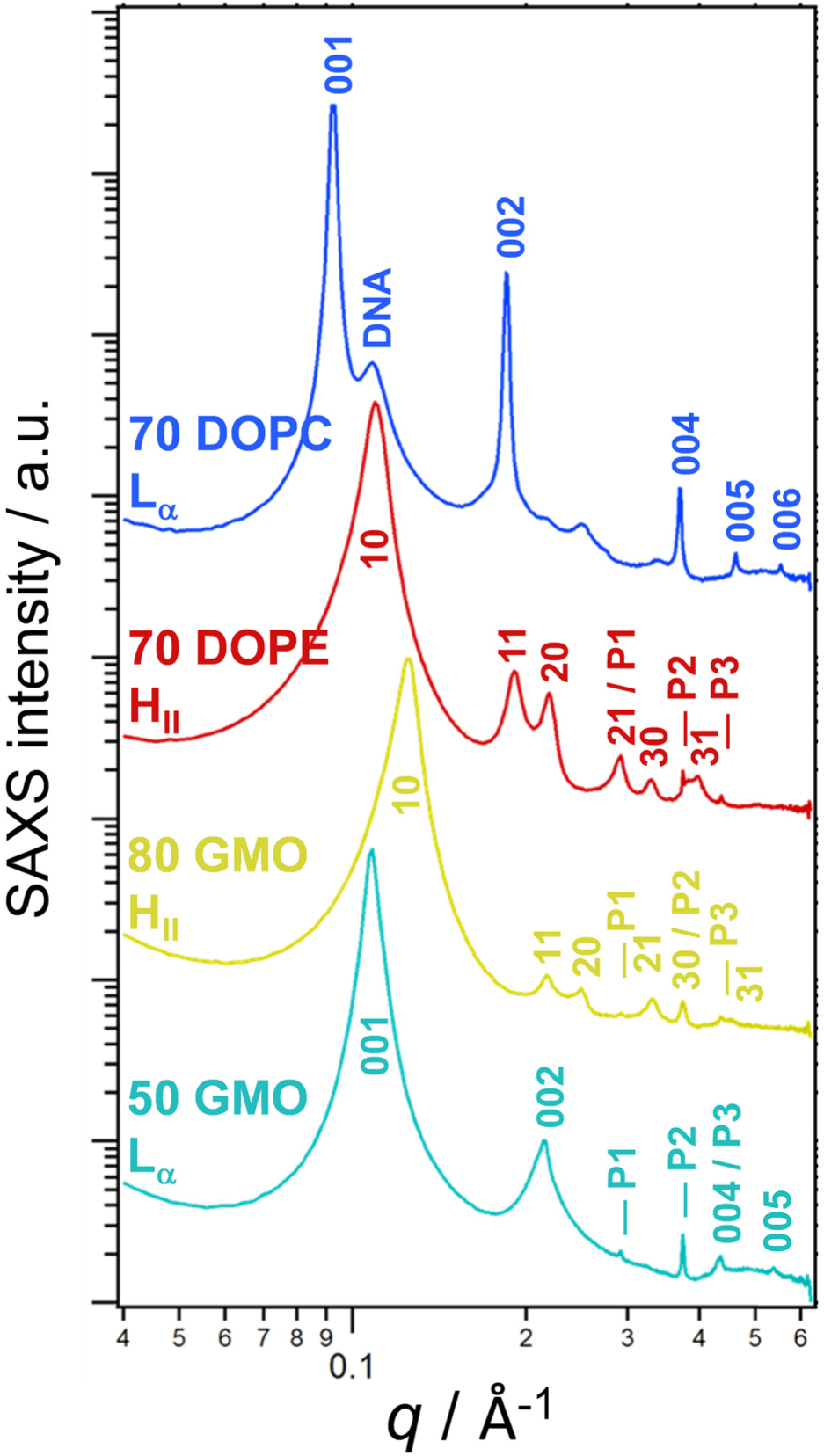
SAXS intensity profiles and analysis of PTX-loaded lipoplexes prepared from binary lipid mixtures of cationic DOTAP with neutral DOPC, DOPE, or GMO. Indexing of the scattering peaks reveals the self-assembled structures of the lipoplexes. Peak assignments are indicated for the L_α_^C^ (00L) and H ^C^ (HK) membrane structures, the DNA–DNA interaxial spacing of the L_α_^C^ phase (DNA), and PTX crystals (P1, P2, P3). Lipid mixtures contained 2 mol% PTX, neutral lipid at the indicated mol% and the remainder DOTAP. The lipid mixtures were complexed with calf thymus DNA at a lipid:DNA charge ratio of 1:1.

**Fig. 6.**
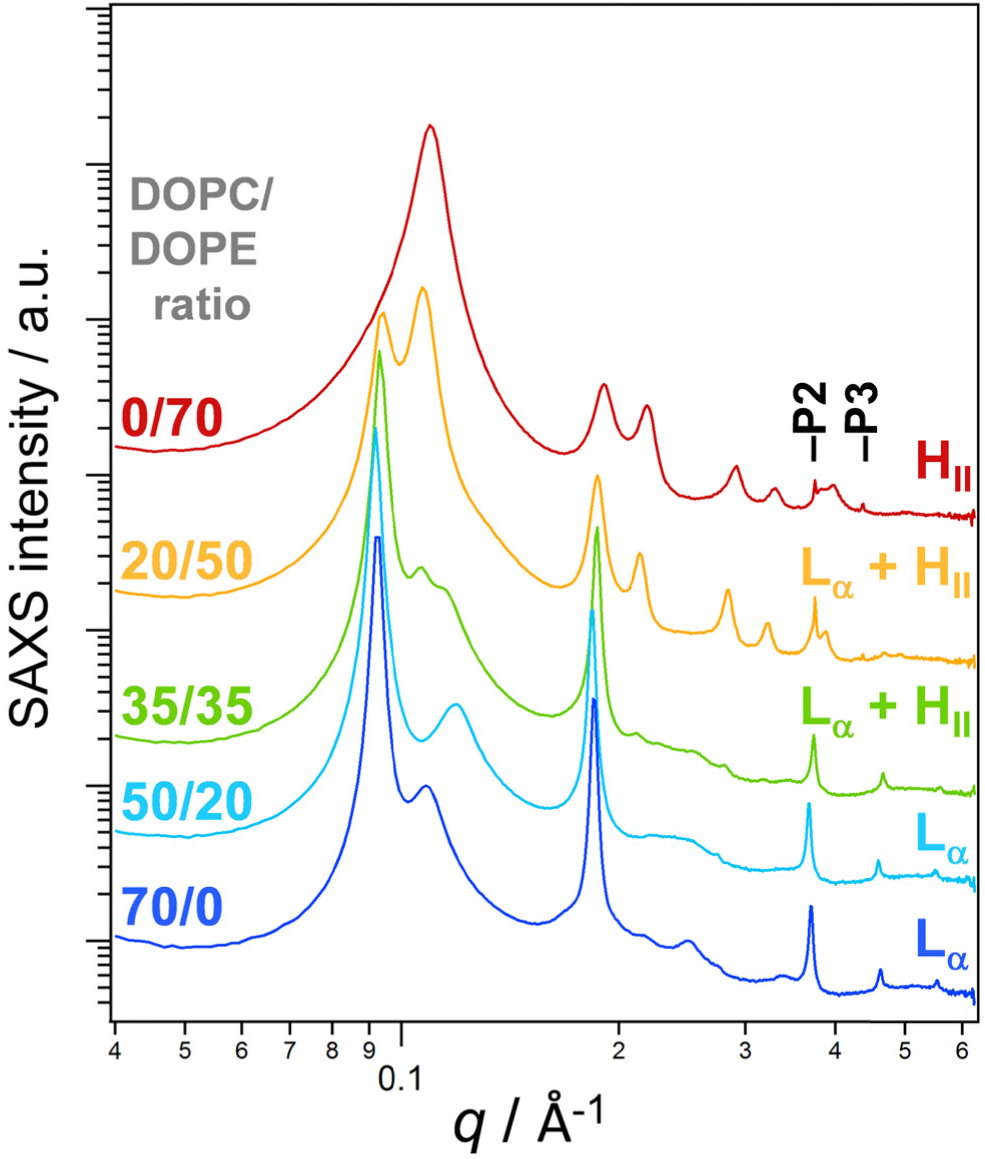
SAXS intensity profiles and analysis of PTX-loaded lipoplexes prepared from lipid mixtures of DOTAP with varied amounts of DOPC and DOPE. The DOPC:DOPE ratio decreases from top to bottom orange; 35:35 (0:70 in in green; red; 20:50 in 50:20 in light blue; 70:0 in dark blue). Lipoplex structures (L_α_^C^ and/or H ^C^) were determined from the structure factor and are reported on the right of the intensity profiles. The peaks labeled P2 and P3 result from PTX crystals present in the with high DOPE content. Lipid mixtures contained 2 DOTAP, and mol% PTX, 28 mol% DOPC/DOPE as indicated for a total of 70 mol%. The lipid mixtures were complexed with calf thymus DNA at a lipid:DNA charge ratio of 1:1.

The SAXS data of lipoplexes formed from ternary lipid mixtures of DOTAP/DOPC/DOPE with DOPE (negative *C*_0_) content between 0 and 70 mol% is shown in Fig. 6. As already shown in Fig. 5, the formulations with 70 mol% DOPC (dark blue) and 70 mol% DOPE (red) yield the L_α_^C^ and H ^C^ lipoplex phases, respectively. The formul**a**tion containing a DOPC/DOPE mixture at a ratio of 20/50 (yellow) yields a coexisting mixture of L_α_^C^ and H_II_^C^ lipoplexes. When the amount of DOPC in the lipid formulation is increased to equal amounts of DOPC and DOPE (green, 35/35 ratio), the L_α_^C^ phase dominates, but w**e** also observe a minor fraction of H_II_^C^ lipoplexes. Further increasing DOPC content to 50 mol% (light blue, 20 mol% DOPE) eliminates the H_II_^C^ phase, and only the L_α_^C^ phase is observed.

Combining the information obtained by SAXS and the kinetic phase diagrams shows that PTX solubility is lower in formulations of lipids with negative spontaneous curvature even when their DNA complexes exhibit L ^C^ morphology. This point is best illustrated by the formulations containing 50 mol% GMO sample and a mixture of DOPC and **D**OPE at a 50:20 ratio. Both formulations form DNA complexes in the L_α_^C^ phase but have decreased PTX solubility compared to the formulation at 70 mol% DOPC which lacks lipids with negative *C*_0_ (and also forms the L_α_^C^ phase).

## Discussion and Conclusion

We investigated lipids promoting a negative spontaneous curvature (*C*_0_ < 0) with regard to their suitability as carriers for the cancer chemothera**p**y drug PTX, because such lipids have shown beneficial effects as carriers of nucleic acids (in**c**luding in current mRNA vaccines). Our goal was further to correlate the findings to the physicochemical properties of the lipids. Intriguingly, when used as the neutral lipid in cationic lipid-base**d** formulations of PTX, both DOPE and GMO (*C*_0_ < 0) showed significantly reduced PTX solubility compared to DOPC (*C*_0_ = 0). Crystallization of membrane-solubilized PTX occurred particularly rapidly in formulations containing GMO. It is likely that this drastic decrease in PTX solubility is in part due to molecular structure: GMO has only one acyl tail whereas DOPE (as well as DOPC) has two, and therefore GMO contributes half the amount of hydrophobic chains to associate with and solubilize PTX. This likely reduces the area and volume of the hydrophobic membrane domain where PTX resides. However, the reduced number of acyl tails is insufficient to explain the low solubility of PTX in GMO-containing formulations. This is evident if we compare formulations containing 80 mol% GMO and 1 mol% PTX, 50 mol% GMO and 1.5 mol% PTX, 70 mol% DOPE and 2 mol% PTX, and 70 mol% DOPC and 2 mol% GMO. These contain 118 oleyl tails per PTX molecule for the formulation with 80 mol% GMO and 98 oleyl tails per PTX molecule for the three others. However, the time to crystallization recorded in the kinetic phase diagrams increases for these formulations from 4 h (GMO-containing formulations) to 8 h (DOPE) to >20 days (DOPC). This strongly suggests that there are additional factors promoting PTX phase separation and crystal nucleation and growth in the presence of GMO. For example, unlike cholesterol, incorporation of which is known to result in tight membranes with a high permeability barrier, single-tailed GMO may contribute to a lowering of the permeability barrier. This would allow water molecules to diffuse more readily across the membrane, leading to PTX phase separation.

To determine whether the lyotropic morphology of the formulations is one of these factors, we performed SAXS experiments on complexes of the cationic liposome formulations with DNA. While formulations with a high content of GMO or DOPE formed the H_II_^C^ phase lipoplexes expected for lipids promoting *C*_0_ < 0, we also identified formulations of both lipids that yielded complexes in the lamellar L_α_^C^ phase. These formulations nonetheless exhibited poorer PTX solubility than the formulation containing only DOPC as the neutral lipid. This suggests that local intermolecular interactions between the lipids and PTX exist which affect PTX solubility more than the self-assembled structures of the lipids.

In conclusion, we have found that DOPE and GMO, in contrast to their role in the delivery of nucleic acids, are not suitable lipids for the delivery of PTX (and likely other hydrophobic drugs) because they promote PTX phase separation from the lipid-based carrier. There likely is a distinct intermolecular contribution to this limited PTX solubility which warrants further investigation. In addition to studies with other lipids, we anticipate that molecular dynamics simulations may prove useful for probing how intermolecular interactions either promote PTX phase separation or improve its solubility. The current study paves the way for future studies of PTX solubility in liposomes containing positive spontaneous curvature (*C*_0_ > 0) inducing lipids, including lipids which contain large multivalent headgroups [75,83,91-97].

## Acknowledgements

The research was supported by the National Institutes of Health under award R01GM130769 (mechanistic studies on developing lipid nanoparticles for drug delivery). Partial support was provided by the National Science Foundation under award DMR-1807327 (membrane phase behavior of lipid nanoparticles). V.M.S. was supported by the National Science Foundation Graduate Research Fellowship Program under Grant No. DGE 1144085. C.R.S. acknowledges sabbatical leave for Spring quarter 2023, which allowed for scholarly activities related to the research reported here.

## Author contributions

The experiments were conducted by VS, SM, JC, and MM. The data were analyzed and discussed by VS, KKE, YL, and CRS. The first draft of the manuscript was written by VS and KKE; CRS and YL contributed to the final draft, which was reviewed by all authors.

## References

1. Wani, M. C.; Taylor, H. L.; Wall, M. E.; Coggon, P.; McPhail, A. T.: Plant antitumor agents. VI. Isolation and structure of taxol, a novel antileukemic and antitumor agent from Taxus brevifolia. J. Am. Chem. Soc. 1971, 93, 2325–2327. DOI: 10.1021/ja00738a045.

2. Rowinsky, E. K.; Donehower, R. C.: Paclitaxel (Taxol). N. Engl. J. Med. 1995, 332, 1004–1014. DOI: 10.1056/nejm199504133321507.

3. Markman, M.; Mekhail, T. M.: Paclitaxel in cancer therapy. Expert Opin. Pharmacother. 2002, 3, 755–766. DOI: 10.1517/14656566.3.6.755.

4. Ramalingam, S.; Belani, C. P.: Paclitaxel for non-small cell lung cancer. Expert Opin. Pharmacother. 2004, 5, 1771–1780. DOI: 10.1517/14656566.5.8.1771.

5. Hironaka, S.; Zenda, S.; Boku, N.; Fukutomi, A.; Yoshino, T.; Onozawa, Y.: Weekly paclitaxel as second-line chemotherapy for advanced or recurrent gastric cancer. Gastric Cancer 2006, 9, 14–18. DOI: 10.1007/s10120-005-0351-6.

6. Sakamoto, J.; Matsui, T.; Kodera, Y.: Paclitaxel chemotherapy for the treatment of gastric cancer. Gastric Cancer 2009, 12, 69–78. DOI: 10.1007/s10120-009-0505-z.

7. Moxley, K. M.; McMeekin, D. S.: Endometrial Carcinoma: A Review of Chemotherapy, Drug Resistance, and the Search for New Agents. Oncologist 2010, 15, 1026–1033. DOI: 10.1634/theoncologist.2010-0087.

8. Bristol-Myers Squibb Company: Taxol® [package insert]. Available at: http://www.accessdata.fda.gov/drugsatfda_docs/label/2011/020262s049lbl.pdf (accessed Oct 04, 2023).

9. Dorr, R. T.: Pharmacology and Toxicology of Cremophor EL Diluent. Ann. Pharmacother. 1994, 28, S11–S14. DOI: 10.1177/10600280940280S503.

10. Weiss, R. B.; Donehower, R. C.; Wiernik, P. H.; Ohnuma, T.; Gralla, R. J.; Trump, D. L.; Baker, J. R., Jr.; Van Echo, D. A.; Von Hoff, D. D.; Leyland-Jones, B.: Hypersensitivity reactions from taxol. J. Clin. Oncol. 1990, 8, 1263–1268. DOI: 10.1200/jco.1990.8.7.1263.

11. Gelderblom, H.; Verweij, J.; Nooter, K.; Sparreboom, A.: Cremophor EL: the drawbacks and advantages of vehicle selection for drug formulation. Eur. J. Cancer 2001, 37, 1590–1598. DOI: 10.1016/S0959-8049(01)00171-X.

12. Bernabeu, E.; Cagel, M.; Lagomarsino, E.; Moretton, M.; Chiappetta, D. A.: Paclitaxel: What has been done and the challenges remain ahead. Int. J. Pharm. (Amsterdam, Neth.) 2017, 526, 474–495. DOI: 10.1016/j.ijpharm.2017.05.016.

13. Sofias, A. M.; Dunne, M.; Storm, G.; Allen, C.: The battle of “nano” paclitaxel. Adv. Drug Delivery Rev. 2017, 122, 20–30. DOI: 10.1016/j.addr.2017.02.003.

14. Dranitsaris, G.; Yu, B.; Wang, L.; Sun, W.; Zhou, Y.; King, J.; Kaura, S.; Zhang, A.; Yuan, P.: Abraxane® versus Taxol® for patients with advanced breast cancer: A prospective time and motion analysis from a Chinese health care perspective. J. Oncol. Pharm. Pract. 2016, 22, 205–211. DOI: 10.1177/1078155214556008.

15. Rugo, H. S.; Barry, W. T.; Moreno-Aspitia, A.; Lyss, A. P.; Cirrincione, C.; Leung, E.; Mayer, E. L.; Naughton, M.; Toppmeyer, D.; Carey, L. A.; Perez, E. A.; Hudis, C.; Winer, E. P.: Randomized Phase III Trial of Paclitaxel Once Per Week Compared With Nanoparticle Albumin-Bound Nab-Paclitaxel Once Per Week or Ixabepilone With Bevacizumab As First-Line Chemotherapy for Locally Recurrent or Metastatic Breast Cancer: CALGB 40502/NCCTG N063H (Alliance). J. Clin. Oncol. 2015, 33, 2361–2369. DOI: 10.1200/jco.2014.59.5298.

16. Gradishar, W. J.; Tjulandin, S.; Davidson, N.; Shaw, H.; Desai, N.; Bhar, P.; Hawkins, M.; O’Shaughnessy, J.: Phase III Trial of Nanoparticle Albumin-Bound Paclitaxel Compared With Polyethylated Castor Oil–Based Paclitaxel in Women With Breast Cancer. J. Clin. Oncol. 2005, 23, 7794–7803. DOI: 10.1200/jco.2005.04.937.

17. Mahtani, R. L.; Parisi, M.; Glück, S.; Ni, Q.; Park, S.; Pelletier, C.; Faria, C.; Braiteh, F.: Comparative effectiveness of early-line nab-paclitaxel vs. paclitaxel in patients with metastatic breast cancer: a US community-based real-world analysis. Cancer Manag. Res. 2018, 10, 249–256. DOI: 10.2147/cmar.s150960.

18. Mustafa, G.; Hassan, D.; Ruiz-Pulido, G.; Pourmadadi, M.; Eshaghi, M. M.; Behzadmehr, R.; Tehrani, F. S.; Rahdar, A.; Medina, D. I.; Pandey, S.: Nanoscale drug delivery systems for cancer therapy using paclitaxel— A review of challenges and latest progressions. Journal of Drug Delivery Science and Technology 2023, 84, 104494. DOI: 10.1016/j.jddst.2023.104494.

19. Koudelka, Š.; Turánek, J.: Liposomal paclitaxel formulations. J. Controlled Release 2012, 163, 322–334. DOI: 10.1016/j.jconrel.2012.09.006.

20. Wang, F.; Porter, M.; Konstantopoulos, A.; Zhang, P.; Cui, H.: Preclinical development of drug delivery systems for paclitaxel-based cancer chemotherapy. J. Controlled Release 2017, 267, 100–118. DOI: 10.1016/j.jconrel.2017.09.026.

21. Rosenblum, D.; Joshi, N.; Tao, W.; Karp, J. M.; Peer, D.: Progress and challenges towards targeted delivery of cancer therapeutics. Nat. Commun. 2018, 9, 1410. DOI: 10.1038/s41467-018-03705-y.

22. Ma, P.; Mumper, R. J.: Paclitaxel nano-delivery systems: a comprehensive review. J. Nanomed. Nanotechnol. 2013, 4, 1000164. DOI: 10.4172/2157-7439.1000164.

23. Tibbitt, M. W.; Dahlman, J. E.; Langer, R.: Emerging Frontiers in Drug Delivery. J. Am. Chem. Soc. 2016, 138, 704–717. DOI: 10.1021/jacs.5b09974.

24. Blanco, E.; Shen, H.; Ferrari, M.: Principles of nanoparticle design for overcoming biological barriers to drug delivery. Nat. Biotechnol. 2015, 33, 941–951. DOI: 10.1038/nbt.3330.

25. Silli, E. K.; Li, M.; Shao, Y.; Zhang, Y.; Hou, G.; Du, J.; Liang, J.; Wang, Y.: Liposomal nanostructures for Gemcitabine and Paclitaxel delivery in pancreatic cancer. Eur. J. Pharm. Biopharm. 2023, 192, 13–24. DOI: 10.1016/j.ejpb.2023.09.014.

26. Teo, P. Y.; Cheng, W.; Hedrick, J. L.; Yang, Y. Y.: Co-delivery of drugs and plasmid DNA for cancer therapy. Adv. Drug Delivery Rev. 2016, 98, 41–63. DOI: 10.1016/j.addr.2015.10.014.

27. Fasol, U.; Frost, A.; Büchert, M.; Arends, J.; Fiedler, U.; Scharr, D.; Scheuenpflug, J.; Mross, K.: Vascular and pharmacokinetic effects of EndoTAG-1 in patients with advanced cancer and liver metastasis. Ann. Oncol. 2012, 23, 1030–1036. DOI: 10.1093/annonc/mdr300.

28. Campbell, R. B.; Ying, B.; Kuesters, G. M.; Hemphill, R.: Fighting cancer: From the bench to bedside using second generation cationic liposomal therapeutics. J. Pharm. Sci. 2009, 98, 411–429. DOI: 10.1002/jps.21458.

29. Strieth, S.; Eichhorn, M. E.; Sauer, B.; Schulze, B.; Teifel, M.; Michaelis, U.; Dellian, M.: Neovascular targeting chemotherapy: Encapsulation of paclitaxel in cationic liposomes impairs functional tumor microvasculature. Int. J. Cancer 2004, 110, 117–124. DOI: 10.1002/ijc.20083.

30. Strieth, S.; Nussbaum, C. F.; Eichhorn, M. E.; Fuhrmann, M.; Teifel, M.; Michaelis, U.; Berghaus, A.; Dellian, M.: Tumor-selective vessel occlusions by platelets after vascular targeting chemotherapy using paclitaxel encapsulated in cationic liposomes. Int. J. Cancer 2008, 122, 452–460. DOI: 10.1002/ijc.23088.

31. Kunstfeld, R.; Wickenhauser, G.; Michaelis, U.; Teifel, M.; Umek, W.; Naujoks, K.; Wolff, K.; Petzelbauer, P.: Paclitaxel Encapsulated in Cationic Liposomes Diminishes Tumor Angiogenesis and Melanoma Growth in a “Humanized” SCID Mouse Model. J. Invest. Dermatol. 2003, 120, 476–482. DOI: 10.1046/j.1523-1747.2003.12057.x.

32. Schmitt-Sody, M.; Strieth, S.; Krasnici, S.; Sauer, B.; Schulze, B.; Teifel, M.; Michaelis, U.; Naujoks, K.; Dellian, M.: Neovascular Targeting Therapy: Paclitaxel Encapsulated in Cationic Liposomes Improves Antitumoral Efficacy. Clin. Cancer Res. 2003, 9, 2335–2341.

33. Sercombe, L.; Veerati, T.; Moheimani, F.; Wu, S. Y.; Sood, A. K.; Hua, S.: Advances and Challenges of Liposome Assisted Drug Delivery. Front. Pharmacol. 2015, 6, 286. DOI: 10.3389/fphar.2015.00286.

34. Barenholz, Y.: Design of liposome-based drug carriers: from basic research to application as approved drugs. In Medical applications of liposomes, Lasic, D. D.; Papahadjopoulos, D., Eds. Elsevier: Amsterdam, 1998; pp. 545–565. DOI: 10.1016/B978-044482917-7/50031-4.

35. Ewert, K. K.; Scodeller, P.; Simón-Gracia, L.; Steffes, V. M.; Wonder, E. A.; Teesalu, T.; Safinya, C. R.: Cationic Liposomes as Vectors for Nucleic Acid and Hydrophobic Drug Therapeutics. Pharmaceutics 2021, 13, 1365. PMCID: PMC8465808. DOI: 10.3390/pharmaceutics13091365.

36. Kulkarni, J. A.; Witzigmann, D.; Thomson, S. B.; Chen, S.; Leavitt, B. R.; Cullis, P. R.; van der Meel, R.: The current landscape of nucleic acid therapeutics. Nat. Nanotechnol. 2021. DOI: 10.1038/s41565-021-00898-0.

37. Witzigmann, D.; Kulkarni, J. A.; Leung, J.; Chen, S.; Cullis, P. R.; van der Meel, R.: Lipid nanoparticle technology for therapeutic gene regulation in the liver. Adv. Drug Delivery Rev. 2020, 159, 344–363. DOI: 10.1016/j.addr.2020.06.026.

38. Kulkarni, J. A.; Witzigmann, D.; Chen, S.; Cullis, P. R.; van der Meel, R.: Lipid Nanoparticle Technology for Clinical Translation of siRNA Therapeutics. Acc. Chem. Res. 2019, 52, 2435–2444. DOI: 10.1021/acs.accounts.9b00368.

39. Steffes, V. M.; Murali, M. M.; Park, Y.; Fletcher, B. J.; Ewert, K. K.; Safinya, C. R.: Distinct Solubility and Cytotoxicity Regimes of Paclitaxel-Loaded Cationic Liposomes at Low and High Drug Content Revealed by Kinetic Phase Behavior and Cancer Cell Viability Studies. Biomaterials 2017, 145, 242–255. PMCID: PMC5610109. DOI: 10.1016/j.biomaterials.2017.08.026.

40. Campbell, R. B.; Balasubramanian, S. V.; Straubinger, R. M.: Influence of cationic lipids on the stability and membrane properties of paclitaxel-containing liposomes. J. Pharm. Sci. 2001, 90, 1091–1105. DOI: 10.1002/jps.1063.

41. Bernsdorff, C.; Reszka, R.; Winter, R.: Interaction of the anticancer agent Taxol™ (paclitaxel) with phospholipid bilayers. J. Biomed. Mater. Res. 1999, 46, 141–149. DOI: 10.1002/(sici)1097-4636(199908)46:2<141::aid-jbm2>3.0.co;2-u.

42. Barenholz, Y.: Doxil® — The first FDA-approved nano-drug: Lessons learned. J. Controlled Release 2012, 160, 117–134. DOI: 10.1016/j.jconrel.2012.03.020.

43. Fritze, A.; Hens, F.; Kimpfler, A.; Schubert, R.; Peschka-Süss, R.: Remote loading of doxorubicin into liposomes driven by a transmembrane phosphate gradient. Biochim. Biophys. Acta, Biomembr. 2006, 1758, 1633–1640. DOI: 10.1016/j.bbamem.2006.05.028.

44. Li, X.; Hirsh, D. J.; Cabral-Lilly, D.; Zirkel, A.; Gruner, S. M.; Janoff, A. S.; Perkins, W. R.: Doxorubicin physical state in solution and inside liposomes loaded via a pH gradient. Biochim. Biophys. Acta, Biomembr. 1998, 1415, 23–40. DOI: 10.1016/S0005-2736(98)00175-8.

45. Weaver, B. A.: How Taxol/paclitaxel kills cancer cells. Mol. Biol. Cell 2014, 25, 2677–2681. DOI: 10.1091/mbc.E14-04-0916.

46. Schiff, P. B.; Fant, J.; Horwitz, S. B.: Promotion of microtubule assembly in vitro by taxol. Nature 1979, 277, 665–667. DOI: 10.1038/277665a0.

47. Jordan, M. A.; Wendell, K.; Gardiner, S.; Brent Derry, W.; Copp, H.; Wilson, L.: Mitotic Block Induced in HeLa Cells by Low Concentrations of Paclitaxel (Taxol) Results in Abnormal Mitotic Exit and Apoptotic Cell Death. Cancer Res. 1996, 56, 816–825.

48. Jordan, M. A.; Wilson, L.: Microtubules as a target for anticancer drugs. Nat. Rev. Cancer 2004, 4, 253–265. DOI: 10.1038/nrc1317.

49. Jordan, M. A.; Toso, R. J.; Thrower, D.; Wilson, L.: Mechanism of mitotic block and inhibition of cell proliferation by taxol at low concentrations. Proc. Natl. Acad. Sci. U. S. A. 1993, 90, 9552–9556. DOI: 10.1073/pnas.90.20.9552.

50. Yvon, A.-M. C.; Wadsworth, P.; Jordan, M. A.: Taxol Suppresses Dynamics of Individual Microtubules in Living Human Tumor Cells. Mol. Biol. Cell 1999, 10, 947–959. DOI: 10.1091/mbc.10.4.947.

51. Steffes, V. M.; Zhang, Z.; Ewert, K. K.; Safinya, C. R.: The Influence of Multivalent Charge and PEGylation on Shape Transitions in Fluid Lipid Assemblies: From Vesicles to Discs, Rods, and Spheres. bioRxiv 2023, 2023.08.09.552538. DOI: 10.1101/2023.08.09.552538.

52. Simón-Gracia, L.; Scodeller, P.; Fisher, W. S.; Sidorenko, V.; Steffes, V. M.; Ewert, K. K.; Safinya, C. R.; Teesalu, T.: Paclitaxel-Loaded Cationic Fluid Lipid Nanodiscs and Liposomes with Brush-Conformation PEG Chains Penetrate Breast Tumors and Trigger Caspase-3 Activation. ACS Appl. Mater. Interfaces 2022, 14, 56613–56622. DOI: 10.1021/acsami.2c17961.

53. Zhen, Y.; Ewert, K. K.; Fisher, W. S.; Steffes, V. M.; Li, Y.; Safinya, C. R.: Paclitaxel loading in cationic liposome vectors is enhanced by replacement of oleoyl with linoleoyl tails with distinct lipid shapes. Sci. Rep. 2021, 11, 7311. PMCID: PMC8012651. DOI: 10.1038/s41598-021-86484-9.

54. Steffes, V. M.; Zhang, Z.; MacDonald, S.; Crowe, J.; Ewert, K. K.; Carragher, B.; Potter, C. S.; Safinya, C. R.: PEGylation of Paclitaxel-Loaded Cationic Liposomes Drives Steric Stabilization of Bicelles and Vesicles thereby Enhancing Delivery and Cytotoxicity to Human Cancer Cells. ACS Appl. Mater. Interfaces 2020, 12, 151–162. PMCID: PMC6984750. DOI: 10.1021/acsami.9b16150.

55. Hoy, S. M.: Patisiran: First Global Approval. Drugs 2018, 78, 1625–1631. DOI: 10.1007/s40265-018-0983-6.

56. Polack, F. P.; Thomas, S. J.; Kitchin, N.; Absalon, J.; Gurtman, A.; Lockhart, S.; Perez, J. L.; Pérez Marc, G.; Moreira, E. D.; Zerbini, C.; Bailey, R.; Swanson, K. A.; Roychoudhury, S.; Koury, K.; Li, P.; Kalina, W. V.; Cooper, D.; Frenck, R. W.; Hammitt, L. L.; Türeci, Ö.; Nell, H.; Schaefer, A.; Ünal, S.; Tresnan, D. B.; Mather, S.; Dormitzer, P. R.; Şahin, U.; Jansen, K. U.; Gruber, W. C.: Safety and Efficacy of the BNT162b2 mRNA Covid-19 Vaccine. N. Engl. J. Med. 2020, 383, 2603–2615. DOI: 10.1056/NEJMoa2034577.

57. United States Food and Drug Administration: Pfizer-BioNTech COVID-19 Vaccine Emergency Use Authorization Fact Sheet for Healthcare Providers Administering Vaccine. Available at: https://www.fda.gov/media/144413/download (accessed 10 June 2021).

58. Baden, L. R.; El Sahly, H. M.; Essink, B.; Kotloff, K.; Frey, S.; Novak, R.; Diemert, D.; Spector, S. A.; Rouphael, N.; Creech, C. B.; McGettigan, J.; Khetan, S.; Segall, N.; Solis, J.; Brosz, A.; Fierro, C.; Schwartz, H.; Neuzil, K.; Corey, L.; Gilbert, P.; Janes, H.; Follmann, D.; Marovich, M.; Mascola, J.; Polakowski, L.; Ledgerwood, J.; Graham, B. S.; Bennett, H.; Pajon, R.; Knightly, C.; Leav, B.; Deng, W.; Zhou, H.; Han, S.; Ivarsson, M.; Miller, J.; Zaks, T.: Efficacy and Safety of the mRNA-1273 SARS-CoV-2 Vaccine. N. Engl. J. Med. 2021, 384, 403–416. DOI: 10.1056/NEJMoa2035389.

59. Corbett, K. S.; Edwards, D.; Leist, S. R.; Abiona, O. M.; Boyoglu-Barnum, S.; Gillespie, R. A.; Himansu, S.; Schäfer, A.; Ziwawo, C. T.; DiPiazza, A. T.; Dinnon, K. H.; Elbashir, S. M.; Shaw, C. A.; Woods, A.; Fritch, E. J.; Martinez, D. R.; Bock, K. W.; Minai, M.; Nagata, B. M.; Hutchinson, G. B.; Bahl, K.; Garcia-Dominguez, D.; Ma, L.; Renzi, I.; Kong, W.-P.; Schmidt, S. D.; Wang, L.; Zhang, Y.; Stevens, L. J.; Phung, E.; Chang, L. A.; Loomis, R. J.; Altaras, N. E.; Narayanan, E.; Metkar, M.; Presnyak, V.; Liu, C.; Louder, M. K.; Shi, W.; Leung, K.; Yang, E. S.; West, A.; Gully, K. L.; Wang, N.; Wrapp, D.; Doria-Rose, N. A.; Stewart-Jones, G.; Bennett, H.; Nason, M. C.; Ruckwardt, T. J.; McLellan, J. S.; Denison, M. R.; Chappell, J. D.; Moore, I. N.; Morabito, K. M.; Mascola, J. R.; Baric, R. S.; Carfi, A.; Graham, B. S.: SARS-CoV-2 mRNA Vaccine Development Enabled by Prototype Pathogen Preparedness. bioRxiv 2020, 2020.06.11.145920. DOI: 10.1101/2020.06.11.145920.

60. Krasnici, S.; Werner, A.; Eichhorn, M. E.; Schmitt-Sody, M.; Pahernik, S. A.; Sauer, B.; Schulze, B.; Teifel, M.; Michaelis, U.; Naujoks, K.; Dellian, M.: Effect of the surface charge of liposomes on their uptake by angiogenic tumor vessels. Int. J. Cancer 2003, 105, 561–567. DOI: 10.1002/ijc.11108.

61. Campbell, R. B.; Fukumura, D.; Brown, E. B.; Mazzola, L. M.; Izumi, Y.; Jain, R. K.; Torchilin, V. P.; Munn, L. L.: Cationic Charge Determines the Distribution of Liposomes between the Vascular and Extravascular Compartments of Tumors. Cancer Res. 2002, 62, 6831–6836.

62. Eichhorn, M. E.; Ischenko, I.; Luedemann, S.; Strieth, S.; Papyan, A.; Werner, A.; Bohnenkamp, H.; Guenzi, E.; Preissler, G.; Michaelis, U.; Jauch, K.-W.; Bruns, C. J.; Dellian, M.: Vascular targeting by EndoTAG™-1 enhances therapeutic efficacy of conventional chemotherapy in lung and pancreatic cancer. Int. J. Cancer 2010, 126, 1235–1245. DOI: 10.1002/ijc.24846.

63. Thurston, G.; McLean, J. W.; Rizen, M.; Baluk, P.; Haskell, A.; Murphy, T. J.; Hanahan, D.; McDonald, D. M.: Cationic liposomes target angiogenic endothelial cells in tumors and chronic inflammation in mice. J. Clin. Invest. 1998, 101, 1401–1413. DOI: 10.1172/jci965.

64. Bode, C.; Trojan, L.; Weiss, C.; Kraenzlin, B.; Michaelis, U.; Teifel, M.; Alken, P.; Michel, M. S.: Paclitaxel encapsulated in cationic liposomes: A new option for neovascular targeting for the treatment of prostate cancer. Oncol. Rep. 2009, 22, 321–326. DOI: 10.3892/or_00000440.

65. Ho, E. A.; Ramsay, E.; Ginj, M.; Anantha, M.; Bregman, I.; Sy, J.; Woo, J.; Osooly-Talesh, M.; Yapp, D. T.; Bally, M. B.: Characterization of Cationic Liposome Formulations Designed to Exhibit Extended Plasma Residence Times and Tumor Vasculature Targeting Properties. J. Pharm. Sci. 2010, 99, 2839–2853. DOI: 10.1002/jps.22043.

66. Wonder, E.; Simón-Gracia, L.; Scodeller, P.; Majzoub, R. N.; Kotamraju, V. R.; Ewert, K. K.; Teesalu, T.; Safinya, C. R.: Competition of charge-mediated and specific binding by peptide-tagged cationic liposome–DNA nanoparticles in vitro and in vivo. Biomaterials 2018, 166, 52–63. PMCID: PMC5944340. DOI: 10.1016/j.biomaterials.2018.02.052.

67. Fetterly, G. J.; Straubinger, R. M.: Pharmacokinetics of paclitaxel-containing liposomes in rats. AAPS PharmSci 2003, 5, E32. DOI: 10.1208/ps050432.

68. Sharma, A.; Sharma, U. S.; Straubinger, R. M.: Paclitaxel-liposomes for intracavitary therapy of intraperitoneal P388 leukemia. Cancer Lett. 1996, 107, 265–272. DOI: 10.1016/0304-3835(96)04380-7.

69. Briuglia, M.-L.; Rotella, C.; McFarlane, A.; Lamprou, D. A.: Influence of cholesterol on liposome stability and on in vitro drug release. Drug Delivery Transl. Res. 2015, 5, 231–242. DOI: 10.1007/s13346-015-0220-8.

70. Bedu-Addo, R. K.; Tang, P.; Xu, Y.; Huang, L.: Interaction of Polyethyleneglycol-Phospholipid Conjugates with Cholesterol-Phosphatidylcholine Mixtures: Sterically Stabilized Liposome Formulations. Pharm. Res. 1996, 13, 718–724. DOI: 10.1023/a:1016043431778.

71. Hong, S.-S.; Choi, J. Y.; Kim, J. O.; Lee, M.-K.; Kim, S. H.; Lim, S.-J.: Development of paclitaxel-loaded liposomal nanocarrier stabilized by triglyceride incorporation. Int. J. Nanomed. 2016, 11, 4465–4477. DOI: 10.2147/ijn.s113723.

72. Kannan, V.; Balabathula, P.; Divi, M. K.; Thoma, L. A.; Wood, G. C.: Optimization of drug loading to improve physical stability of paclitaxel-loaded long-circulating liposomes. J. Liposome Res. 2015, 25, 308–315. DOI: 10.3109/08982104.2014.995671.

73. Steffes, V. M. Designing lipid nanoparticles toward targeted drug delivery: Fundamental studies identify key compositional properties to improve formulations for the hydrophobic cancer drug paclitaxel. PhD thesis, University of California, Santa Barbara, 2019.

74. Koltover, I.; Salditt, T.; Rädler, J. O.; Safinya, C. R.: An inverted hexagonal phase of cationic liposome-DNA complexes related to DNA release and delivery. Science 1998, 281, 78–81. DOI: 10.1126/science.281.5373.78.

75. Leal, C.; Ewert, K. K.; Shirazi, R. S.; Bouxsein, N. F.; Safinya, C. R.: Nanogyroids Incorporating Multivalent Lipids: Enhanced Membrane Charge Density and Pore Forming Ability for Gene Silencing. Langmuir 2011, 27, 7691–7697. PMCID: PMC3119580. DOI: 10.1021/la200679x.

76. Leal, C.; Ewert, K. K.; Bouxsein, N. F.; Shirazi, R. S.; Li, Y.; Safinya, C. R.: Stacking of short DNA induces the gyroid cubic-to-inverted hexagonal phase transition in lipid-DNA complexes. Soft Matter 2013, 9, 795–804. PMCID: PMC3587977. DOI: 10.1039/c2sm27018h.

77. Leal, C.; Bouxsein, N. F.; Ewert, K. K.; Safinya, C. R.: Highly Efficient Gene Silencing Activity of siRNA Embedded in a Nanostructured Gyroid Cubic Lipid Matrix. J. Am. Chem. Soc. 2010, 132, 16841–16847. PMCID: PMC2991473. DOI: 10.1021/ja1059763.

78. Godbey, W. T.: Gene Delivery. In An Introduction to Biotechnology, Godbey, W. T., Ed. Woodhead Publishing: 2014; pp. 275–312. DOI: 10.1016/B978-1-907568-28-2.00013-7.

79. Crespo-Barreda, A.; Encabo-Berzosa, M. M.; González-Pastor, R.; Ortíz-Teba, P.; Iglesias, M.; Serrano, J. L.; Martin-Duque, P.: Viral and Nonviral Vectors for In Vivo and Ex Vivo Gene Therapies. In Translating Regenerative Medicine to the Clinic, Laurence, J., Ed. Academic Press: Boston, 2016; pp. 155–177. DOI: 10.1016/B978-0-12-800548-4.00011-5.

80. Rodrigues, L.; Schneider, F.; Zhang, X.; Larsson, E.; Moodie, L. W. K.; Dietz, H.; Papadakis, C. M.; Winter, G.; Lundmark, R.; Hubert, M.: Cellular uptake of self-assembled phytantriol-based hexosomes is independent of major endocytic machineries. J. Colloid Interface Sci. 2019, 553, 820–833. DOI: 10.1016/j.jcis.2019.06.045.

81. Safinya, C. R.; Ewert, K. K.; Majzoub, R. N.; Leal, C.: Cationic liposome-nucleic acid complexes for gene delivery and gene silencing. New J. Chem. 2014, 38, 5164–5172. PMCID: PMC4288823. DOI: 10.1039/c4nj01314j.

82. Bouxsein, N. F.; McAllister, C. S.; Ewert, K. K.; Samuel, C. E.; Safinya, C. R.: Structure and gene silencing activities of monovalent and pentavalent cationic lipid vectors complexed with siRNA. Biochemistry 2007, 46, 4785–4792. DOI: 10.1021/bi062138l.

83. Ahmad, A.; Evans, H. M.; Ewert, K.; George, C. X.; Samuel, C. E.; Safinya, C. R.: New multivalent cationic lipids reveal bell curve for transfection efficiency versus membrane charge density: Lipid-DNA complexes for gene delivery. J. Gene. Med. 2005, 7, 739–748. DOI: 10.1002/jgm.717.

84. Lin, A. J.; Slack, N. L.; Ahmad, A.; George, C. X.; Samuel, C. E.; Safinya, C. R.: Three-dimensional imaging of lipid gene-carriers: Membrane charge density controls universal transfection behavior in lamellar cationic liposome-DNA complexes. Biophys. J. 2003, 84, 3307–3316. DOI: 10.1016/S0006-3495(03)70055-1.

85. Kulkarni, C. V.; Wachter, W.; Iglesias-Salto, G.; Engelskirchen, S.; Ahualli, S.: Monoolein: a magic lipid? Phys. Chem. Chem. Phys. 2011, 13, 3004–3021. DOI: 10.1039/c0cp01539c.

86. Larsson, K.: Two cubic phases in monoolein-water system. Nature 1983, 304, 664–664.

87. Briggs, J.; Chung, H.; Caffrey, M.: The Temperature-Composition Phase Diagram and Mesophase Structure Characterization of the Monoolein/Water System. J. Phys. II 1996, 6, 723–751.

88. Leung, S. S. W.; Leal, C.: The stabilization of primitive bicontinuous cubic phases with tunable swelling over a wide composition range. Soft Matter 2019, 15, 1269–1277. DOI: 10.1039/c8sm02059k.

89. Rädler, J. O.; Koltover, I.; Salditt, T.; Safinya, C. R.: Structure of DNA-cationic liposome complexes: DNA intercalation in multilamellar membranes in distinct interhelical packing regimes. Science 1997, 275, 810–814. DOI: 10.1126/science.275.5301.810.

90. Koltover, I.; Salditt, T.; Safinya, C. R.: Phase diagram, stability, and overcharging of lamellar cationic lipid-DNA self-assembled complexes. Biophys. J. 1999, 77, 915–924. PMCID: PMC1300382. DOI: 10.1016/S0006-3495(99)76942-0.

91. Ewert, K.; Ahmad, A.; Evans, H. M.; Schmidt, H.-W.; Safinya, C. R.: Efficient synthesis and cell-transfection properties of a new multivalent cationic lipid for nonviral gene delivery. J. Med. Chem. 2002, 45, 5023–5029. DOI: 10.1021/jm020233w.

92. Ewert, K. K.; Evans, H. M.; Bouxsein, N. F.; Safinya, C. R.: Dendritic cationic lipids with highly charged headgroups for efficient gene delivery. Bioconjugate Chem. 2006, 17, 877–888. DOI: 10.1021/bc050310c.

93. Ewert, K. K.; Evans, H. M.; Zidovska, A.; Bouxsein, N. F.; Ahmad, A.; Safinya, C. R.: A columnar phase of dendritic lipid-based cationic liposome-DNA complexes for gene delivery: Hexagonally ordered cylindrical micelles embedded in a DNA honeycomb lattice. J. Am. Chem. Soc. 2006, 128, 3998–4006. DOI: 10.1021/ja055907h.

94. Farago, O.; Ewert, K.; Ahmad, A.; Evans, H. M.; Grønbech-Jensen, N.; Safinya, C. R.: Transitions between distinct compaction regimes in complexes of multivalent cationic lipids and DNA. Biophys. J. 2008, 95, 836–846. PMCID: PMC2440430. DOI: 10.1529/biophysj.107.124669.

95. Zidovska, A.; Evans, H. M.; Ewert, K. K.; Quispe, J.; Carragher, B.; Potter, C. S.; Safinya, C. R.: Liquid crystalline phases of dendritic lipid-DNA self-assemblies: Lamellar, hexagonal, and DNA bundles. J. Phys. Chem. B 2009, 113, 3694–3703. PMCID: PMC2858692. DOI: 10.1021/jp806863z.

96. Zidovska, A.; Ewert, K. K.; Quispe, J.; Carragher, B.; Potter, C. S.; Safinya, C. R.: Block Liposomes: Vesicles of Charged Lipids with Distinctly Shaped Nanoscale Sphere-, Pear-, Tube-, or Rod-Segments. In Methods in Enzymology: Liposomes, Pt. G, 2009; pp. 111–128. DOI: 10.1016/s0076-6879(09)65006-0.

97. Shirazi, R. S.; Ewert, K. K.; Leal, C.; Majzoub, R. N.; Bouxsein, N. F.; Safinya, C. R.: Synthesis and characterization of degradable multivalent cationic lipids with disulfide-bond spacers for gene delivery. Biochim. Biophys. Acta, Biomembr. 2011, 1808, 2156–66. PMCID: PMC3129426. DOI: 10.1016/j.bbamem.2011.04.020.

